# Gasdermin D is the only Gasdermin that provides non-redundant protection against acute *Salmonella* gut infection

**DOI:** 10.1101/2022.11.24.517575

**Authors:** Stefan A. Fattinger, Luca Maurer, Petra Geiser, Ursina Enz, Suwannee Ganguillet, Ersin Gül, Sanne Kroon, Benjamin Demarco, Markus Furter, Manja Barthel, Pawel Pelczar, Feng Shao, Petr Broz, Mikael E. Sellin, Wolf-Dietrich Hardt

## Abstract

Gasdermins (GSDMs) share a common functional domain structure and are best known for their capacity to form membrane pores. These pores are hallmarks of a specific form of cell death called pyroptosis and mediate the secretion of pro-inflammatory cytokines such as interleukin 1β (IL1β) and interleukin 18 (IL18). Thereby, Gasdermins have been implicated in various immune responses against cancer and infectious diseases such as acute *Salmonella* Typhimurium (*S*.Tm) gut infection. However, to date, we lack a comprehensive functional assessment of the different Gasdermins (GSDMA-E) during *S*.Tm infection *in vivo*. Here, we have performed littermate-controlled oral *S*.Tm infections to investigate the impact of all murine Gasdermins. While GSDMA, -C and -E appear dispensable, we show that GSDMD (i) restricts *S*.Tm loads in the gut tissue and systemic organs, (ii) controls gut inflammation kinetics, and (iii) prevents epithelium disruption by 72h of the infection. Full protection requires GSDMD expression by both bone-marrow-derived lamina propria cells and intestinal epithelial cells (IECs). *In vivo* experiments, 3D- and 2D-enteroid infections further show that infected IEC extrusion proceeds also without GSDMD, but that GSDMD controls the permeabilization and morphology of the extruding cells and affects extrusion kinetics. As such, this work identifies a non-redundant multipronged role of GSDMD in mucosal tissue defence against a common enteric pathogen.

**HIGHLIGHTS:**

- Gasdermin D restricts *Salmonella* Typhimurium (*S*.Tm) translocation across the gut tissue, controls gut inflammation kinetics, and prevents epithelium disruption by 72h of the infection.
- Gasdermins A, C and E appear dispensable for protection against acute *S*.Tm gut infection.
- Gasdermin D in bone-marrow-derived lamina propria cells and intestinal epithelial cells complement each other to suppress gut tissue *S*.Tm loads.
- Gasdermin D is not required for extrusion of infected intestinal epithelial cells but drives their permeabilization and affects qualitative features of the extrusion process.

## INTRODUCTION

Gasdermins make up a protein family including Gasdermin A, B, C, D and E (GSDMA, GSDMB, GSDMC, GSDMD and GSDME, respectively) in humans and GSDMA1-3, GSDMC1-4, GSDMD and GSDME in mice (Broz et al., 2020). All members share a common functional domain structure, in which an inhibitory C-terminal domain is linked to a membrane pore forming N-terminal domain (Broz et al., 2020; Shi et al., 2017). Upon cleavage at the linker region, the N-terminal domain is released to form membrane pores (Aglietti et al., 2016; Ding et al., 2016; Liu et al., 2016; Sborgi et al., 2016). These pores mediate a lytic cell death, which releases inflammatory mediators, such as interleukin 1β (IL1β), interleukin 18 (IL18) and lipids into the extracellular milieu to alert neighboring cells (Kayagaki et al., 2015; Rauch et al., 2017; Shi et al., 2015). Gasdermins can be activated with different efficiency by the cysteine proteases Caspase-1, -3, -4, -8 and -11 and by serine proteases to execute cellular responses, thus mediating immunity against pathogens and cancer (Burgener et al., 2019; Chen et al., 2021; Hou et al., 2020; Kambara et al., 2018; Kayagaki et al., 2015; Liu et al., 2020; Orning et al., 2018; Rogers et al., 2017; Sarhan et al., 2018; Shi et al., 2015; Wang et al., 2017a; Zhang et al., 2020; Zhou et al., 2020; Nozaki et al., 2022; Zhang et al., 2022; Zhao et al., 2022; Lawrence et al., 2022). However, their individual roles, cell type specificities, and possible redundancies during oral bacterial infections, such as those caused by *Salmonella enterica* Serovar Typhimurium (*S*.Tm), have not been comprehensively explored.

*S*.Tm is a major foodborne pathogen, a prevalent cause of diarrheal disease worldwide (Kirk et al., 2015), and a risk factor for inflammatory bowel diseases (Gradel et al., 2009). As shown in streptomycin-pretreated mice - a commonly used mouse model for human *Salmonella* diarrhea - *S*.Tm frequently invades intestinal epithelial cells (IECs) during the acute gut infection, transmigrates into the underlying compartment called the lamina propria, and spreads to systemic organs (Kaiser et al., 2012; Fattinger et al., 2021c). Innate host immune responses against *S*.Tm include the activation of the NAIP/NLRC4 inflammasome, which senses invading *S*.Tm to induce cell death and IL18 release through a mechanism involving Caspase-1 (Fattinger et al., 2021b). During *S*.Tm infection of streptomycin pretreated mice, the NAIP/NLRC4 response particularly in IECs provide a first line of defense, which reduces *S*.Tm loads locally in the gut tissue, as well as restricts pathogen accumulation at systemic sites like the mesenteric lymph nodes (mLN), spleen and liver (Carvalho et al., 2012; Fattinger et al., 2021a; Franchi et al., 2012; Hausmann et al., 2020; Lai et al., 2013; Lara-Tejero et al., 2006; Rauch et al., 2017; Raupach et al., 2006; Sellin et al., 2014; Müller et al., 2016). The IEC’s NAIP/NLRC response limits pathogen spread predominantly by swiftly expelling infected IECs into the gut lumen (Fattinger et al., 2021a; Knodler et al., 2010; Rauch et al., 2017; Sellin et al., 2014). Importantly, epithelial NAIP/NLRC4 not only triggers cell death, but also coordinates the detachment of the infected IEC from the epithelium (a process referred to as IEC extrusion), with a concomitant release of the aforementioned inflammatory mediators (Rauch et al., 2017; Fattinger et al., 2021a; Sellin et al., 2014). In studies using bacteria and/or pure NAIP/NLRC4 ligands, it was shown that GSDMD affects the qualitative features of the IEC extrusion process (Rauch et al., 2017; Samperio Ventayol et al., 2021; Nozaki et al., 2022). However, how and to which extent epithelial GSDMD contributes to the overall defense against *S*.Tm infection *in vivo* remains far from clear. Moreover, we have recently shown that a fraction of the extruding IECs feature activated forms of Caspase-3 and -8 (Fattinger et al., 2021a), which raises the question if other Gasdermins, which can be activated by those caspases, could additionally be involved in IEC extrusion, and thereby contribute to the defense against *S*.Tm. Indeed, recent studies in mice suggested a role for epithelial GSDME in 2,4,6-trinitrobenzenesulfonic acid-induced colitis (Tan et al., 2021), and for epithelial GSDMC in worm-infected mice (Xi et al., 2021; Zhao et al., 2022). Finally, Gasdermins are originally known for their function in phagocytic immune cells. It has been shown that immune cells employ Gasdermins to promote gut inflammation and defense against several gut pathogens *in vivo* (e.g. Chen et al., 2021, 2018a; Ma et al., 2021; Booty and Bryant, 2022; Chauhan et al., 2022; Chen et al., 2018a; Gram et al., 2021; Orning et al., 2018). These observations suggest that not only epithelial Gasdermins, but also Gasdermins expressed by dedicated immune cells may contribute to pathogen restriction during *S*.Tm infection. However, the respective contribution(s) of particular Gasdermins in IECs and immune cells during acute *Salmonella* diarrhea has not yet been systematically addressed.

Here, we have performed a comprehensive assessment of the impact of Gasdermins during oral *S*.Tm infection. Surprisingly, out of all analyzed Gasdermins, only GSDMD appears to significantly protect against *S*.Tm infection, lowering pathogen loads both in the gut mucosal tissue and at systemic organs. We show that epithelial GSDMD impacts how IECs are extruded into the lumen and that GSDMD in IECs and in bone marrow-derived phagocytes of the lamina propria work together to prevent *S*.Tm spread beyond the intestinal barrier.

## RESULTS

### GSDMD deficiency exacerbates disease by 48-72h of *S*.Tm infection

GSDMD is activated by Caspase-1 downstream of inflammasomes such as the NAIP/NLRC4 (Kayagaki et al., 2015; Shi et al., 2015). Streptomycin pretreated mice deficient in the NAIP/NLRC4 inflammasome accumulate higher *S*.Tm loads in the gut tissue as well as in systemic organs and suffer from a TNF-driven collapse of the epithelial barrier by days 2-3 after orogastric infection (Rauch et al., 2017; Fattinger et al., 2021a; Hausmann et al., 2020; Sellin et al., 2014). Therefore, we investigated the effect of GSDMD-deficiency during oral infection with *S*.Tm (SL1344) in streptomycin pretreated mice. In line with previous work (Fujii et al., 2008), the gut mucosa appeared normal in GSDMD-deficient mice and baseline expression levels of inflammatory mediators were indistinguishable from those from matched littermate controls (Fig S1A-C). Upon infection, while gut luminal *S*.Tm colonization was similar across genotypes throughout the infection (Fig S2A), GSDMD-deficiency resulted in 5 to 10-fold elevated systemic *S*.Tm loads at 48h post infection (p.i.), as determined by CFU plating assays of mLN, spleen and liver samples (Fig 1A-C). Of note, CFU levels of *GsdmD*^*+/-*^ mice were comparable to previously observed levels in WT mice indicating that one copy of WT *GsdmD* is sufficient. In both *GsdmD*^*+/-*^ and *GsdmD*^*-/-*^ littermates, we measured high levels of the inflammatory marker lipocalin-2 (LCN2) in the feces at this time point of infection (Fig S2B). Importantly, however, we detected higher *S*.Tm loads in *GsdmD*^*-/-*^ animals specifically in the lamina propria and measured elevated levels of TNF (Fig 1 D-F).

**Figure 1.**
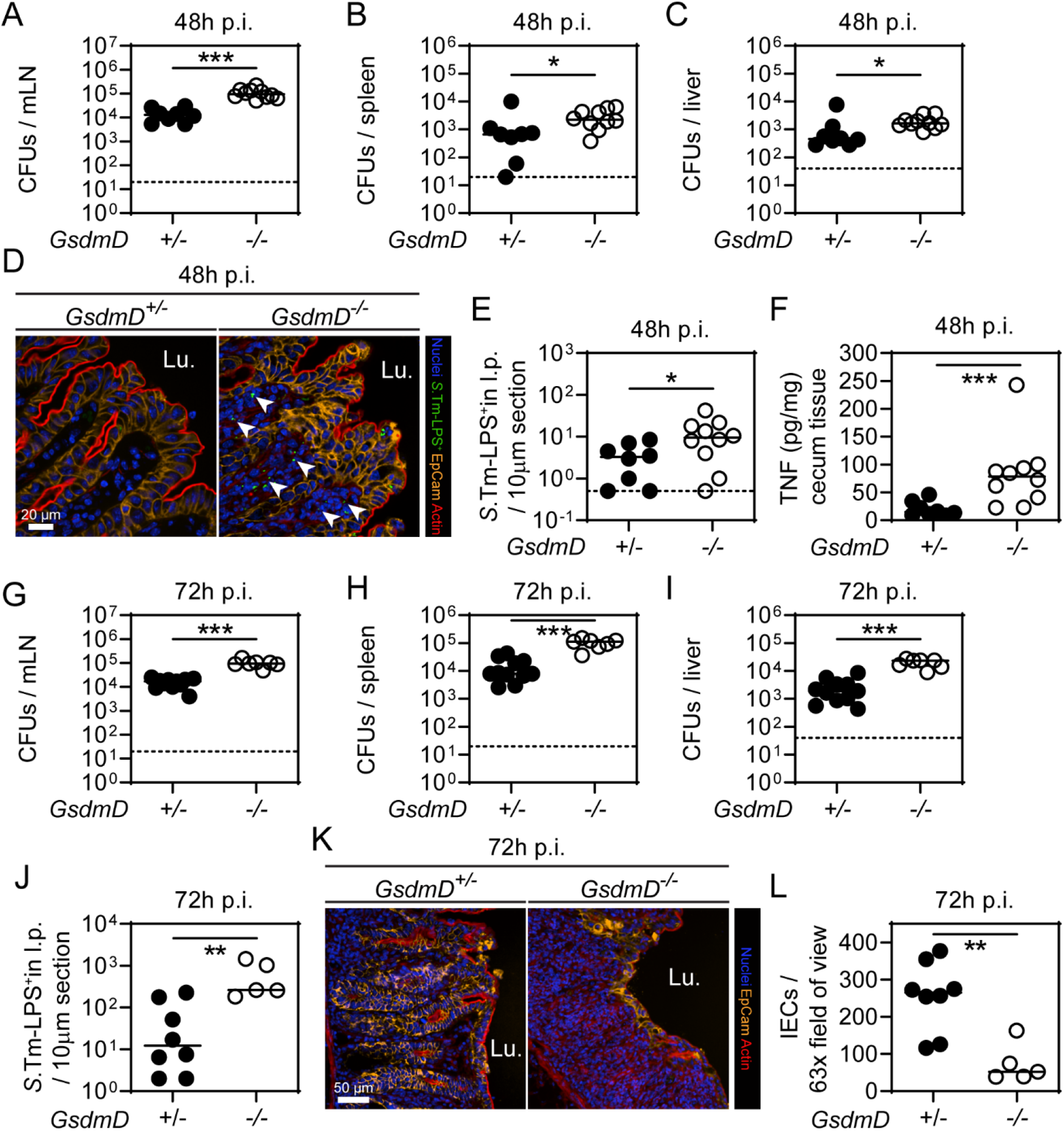
Consequences of GSDMD-deficiency by 48-72h of *S*.Tm infection. (**A-F**) At 48h p.i., GSDMD-deficient mice exhibit elevated *S*.Tm pathogen loads at systemic sides, but also in the gut tissue, leading to high levels of TNF levels compared to heterozygous littermate controls. *S*.Tm pathogen loads in (**A**) mesenteric lymph nodes, (**B**) spleen, and (**C**) liver. (**D**) Representative micrographs of cecum tissue sections, stained for *S*.Tm-LPS. Arrowheads indicate *S*.Tm in lamina propria. Lu. - Lumen. (**E**) Microscopy-based quantification of *S*.Tm-LPS^+^ cells in the lamina propria. (**F**) TNF concentrations in cecum tissue. (**G-L**) At 72h p.i., GSDMD-deficiency still results in elevated *S*.Tm pathogen loads locally and systemically, and epithelial tissue integrity becomes compromised. *S*.Tm CFU pathogen loads in (**G**) mesenteric lymph nodes, (**H**) spleen, and (**I**) liver. (**J**) Microscopy-based quantification of *S*.Tm-LPS^+^ cells in the lamina propria (**K**) Representative micrographs of cecum tissue sections, stained for epithelial marker EpCam. Lu. - Lumen. (**L**) Microscopy-based quantification of IECs per 63x field of view. In A, B, C, E, F, G, H, I, J, L, each data point represents one mouse. ≥5 mice per group from ≥2 independent experiments for each comparison. Line at median. Dotted line represents detection limit. Mann-Whitney U-test (*p<0.05, **p<0.01, ***p<0.001).

As expected in this mouse model, *S*.Tm loads increased from 48 to 72h at systemic sites, but pathogen loads in GSDMD-deficient mice remained consistently higher than in the heterozygous littermate controls (Fig 1G-J, S2C-D). Of further note, the epithelium was severely disrupted by 72h p.i. in *GsdmD*^*-/-*^ mice (Fig 1K). Accordingly, we observed significantly reduced numbers of IECs and more epithelial gaps than in the corresponding control animals at this point of the infection (Fig 1K-L, S2E). This was reminiscent of the day 3 infection phenotypes previously seen in NAIP/NLRC4-deficient mice (Fattinger et al., 2021a). These observations were confirmed by independent experiments using an alternative GSDMD-deficient mouse line (*GsdmD_*fsX^*-/-*^), in which the deficiency was caused by a genetic frameshift instead of a deletion (Fig S2F-Q). Overall, these data suggest that GSDMD is protective against *S*.Tm infection at ∼48-72h p.i., in seeming analogy to NAIP/NLRC4 (Rauch et al., 2017; Fattinger et al., 2021a; Sellin et al., 2014). Both GSDMD and NAIP/NLRC4 limit *S*.Tm loads in the gut tissue, as well as in systemic organs, and prevent the loss of epithelial barrier integrity by 48-72h of infection.

### GSDMD is the only Gasdermin family member with non-redundant protective function in streptomycin pretreated mice infected for 48h with *S*.Tm

Recent studies suggested that Caspase-dependent cell death signaling can be interconnected, potentially leading to the activation of various Gasdermins by various Caspases, and with several Gasdermins proposed to have anti-infection properties (Fattinger et al., 2021b). Since NAIP/NLRC4-driven cell death involves at least Caspase-1 and -8 (Rauch et al., 2017), we explored if other Gasdermins expressed in the gut tissue (Fig S3A) may limit *S*.Tm infection to a similar extent as GSDMD. In particular, we were interested in GSDME since it was shown to be activated by apoptotic Caspases (Orning et al., 2018; Rogers et al., 2017; Sarhan et al., 2018; Chen et al., 2021; Wang et al., 2017b). We infected GSDME-deficient mice for 48h and enumerated *S*.Tm loads in the gut tissue, as well as in systemic organs. No differences in *S*.Tm loads in gut lumen, gut tissue, nor systemic organs, could be detected between GSDME-deficient mice and heterozygous littermate controls (Fig 2A-D, S3B). Furthermore, we investigated if this can be explained by redundancy between GSDMD and GSDME. For this purpose, we compared GSDME proficient and deficient mice in a GSDMD-deficient background (*GsdmD*^*-/-*^*xGsdmE*^*+/-*^ vs. *GsdmD*^*-/-*^*xGsdmE*^*-/-*^). Confirming the results above, the overall *S*.Tm loads in the organs were higher than in GSDMD-proficient mice (compare e.g. Fig 2F with Fig 1A). However, we could still not detect any GSDME-dependent restriction of *S*.Tm, suggesting that GSDME does not influence *S*.Tm loads even in the absence of GSDMD (Fig 2E-H, S3C). Similar observations were made for mice deficient in either all GSDMAs, or all GSDMCs (GSDMA1-3, and GSDMC1-4 respectively). In both mouse lines we could not detect any differences in organ CFU levels by plating assays compared to the respective heterozygous littermate controls (Fig 2I-P, S3D-E). Therefore, out of all described murine Gasdermins, only GSDMD shows a non-redundant function in protecting the gut from *S*.Tm infection in the streptomycin mouse model, as judged by 48h of the infection. Future work could assess if additional Gasdermins may contribute to some specific infection parameter beyond this timeframe. Nevertheless, these data highlight a unique role for GSDMD during acute *S*.Tm gut infection.

**Figure 2.**
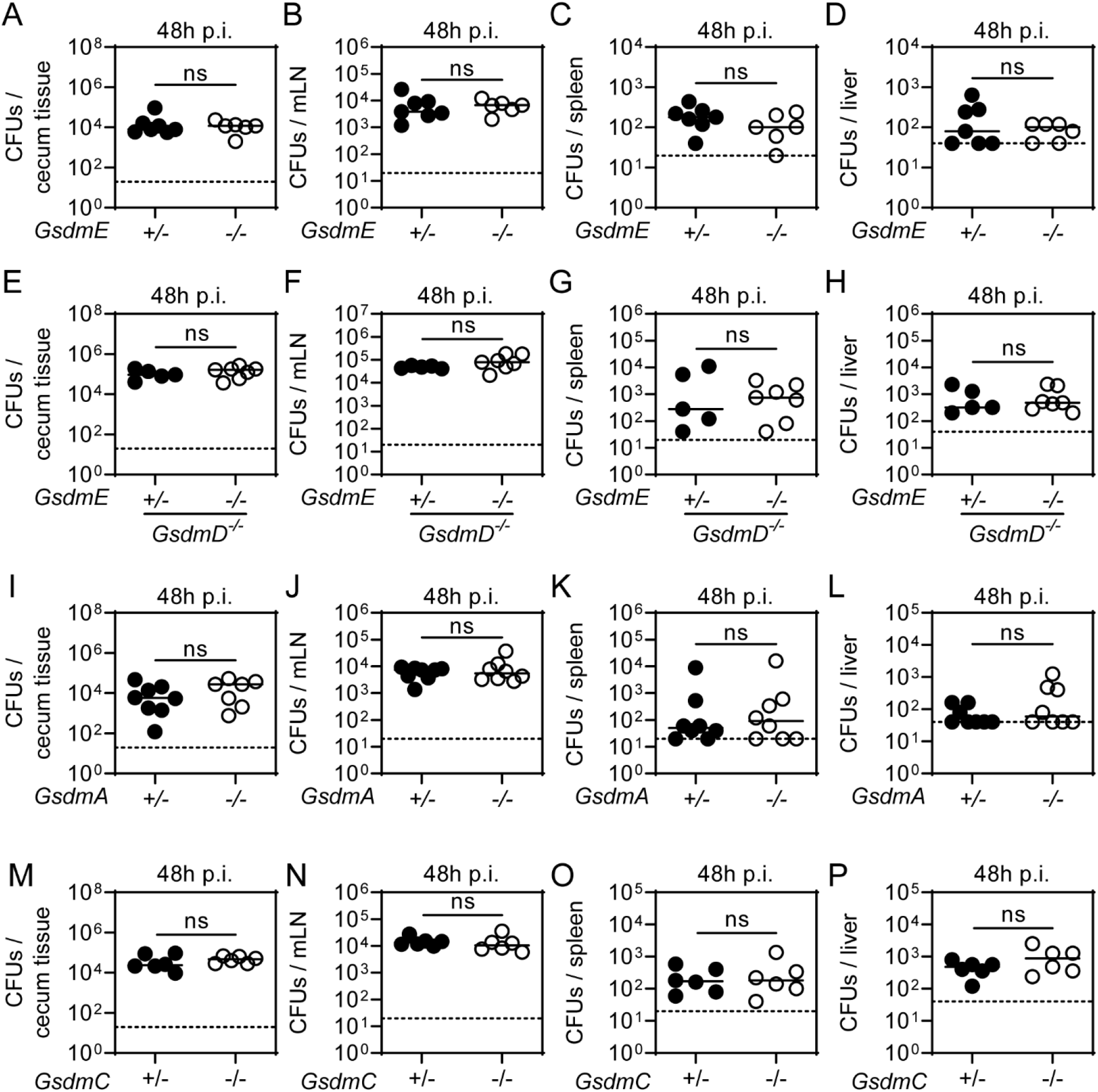
Gasdermin A, C, and E during *S*.Tm infection. (**A-D**) At 48h p.i., GSDME-deficient mice exhibit similar *S*.Tm pathogen loads in the gut tissue and in systemic organs compared to heterozygous littermate controls. *S*.Tm pathogen loads in (**A**) cecum tissue, (**B**) mesenteric lymph nodes, (**C**) spleen, and (**D**) liver. (**E-H**) At 48h p.i., even in the absence of GSDMD, GSDME-deficiency does not result in elevated *S*.Tm pathogen loads in the gut tissue and in systemic organs compared to heterozygous littermate controls. *S*.Tm pathogen loads in (**E**) cecum tissue, (**F**) mesenteric lymph nodes, (**G**) spleen, and (**H**) liver. (**I-L**) At 48h p.i., GSDMA-deficient mice exhibit similar *S*.Tm pathogen loads in the gut tissue and in systemic organs compared to heterozygous littermate controls. *S*.Tm pathogen loads in (**I**) cecum tissue, (**J**) mesenteric lymph nodes, (**K**) spleen, and (**L**) liver. (**M-P**) At 48h p.i., GSDMC-deficient mice exhibit similar *S*.Tm pathogen loads in the gut tissue and in systemic organs compared to heterozygous littermate controls. *S*.Tm pathogen loads in (**M**) cecum tissue, (**N**) mesenteric lymph nodes, (**O**) spleen, and (**P**) liver. Each data point represents one mouse. ≥5 mice per group from ≥2 independent experiments for each comparison. Line at median. Dotted line represents detection limit. Mann-Whitney U-test (ns - not significant).

### Bone marrow derived cells employ GSDMD to restrict *S*.Tm tissue loads

Since GSDMD is known for the induction of pyroptosis in bone marrow (BM) derived macrophages and we observed elevated *S*.Tm loads in the lamina propria of GSDMD-deficient mice, we addressed if GSDMD in BM-derived cells of the lamina propria restricts *S*.Tm *in vivo*. WT mice were gamma-irradiated and reconstituted with BM from either WT or GSDMD-deficient donors, which resulted in >92% transfer efficiency (Fig S4A). When infected, both groups exhibited similar luminal *S*.Tm colonization (Fig S4B). However, *GsdmD*^*-/-*^ BM recipients harbored significantly elevated *S*.Tm loads in the cecum tissue, mLN and spleen at 72h p.i. (Fig 3A-C). Moreover, fluorescence microscopy revealed elevated *S*.Tm loads specifically in the lamina propria compartment (Fig 3D). This demonstrates that lack of GSDMD exclusively in BM derived cells is sufficient to observe higher lamina propria *S*.Tm loads after 72h of infection. Similar observations were made in 48h infections, or in bone marrow chimeric mice derived from GSDMD-deficient recipients, which were infected for 48h or 96h (Fig S4C-D, S4E-H). Accordingly, when *GsdmD*^*-/-*^ mice were infected systemically (intravenous, i.v.), pathogen loads in spleen and liver were again higher than in the heterozygous littermate controls (Fig 3E-F).

**Figure 3.**
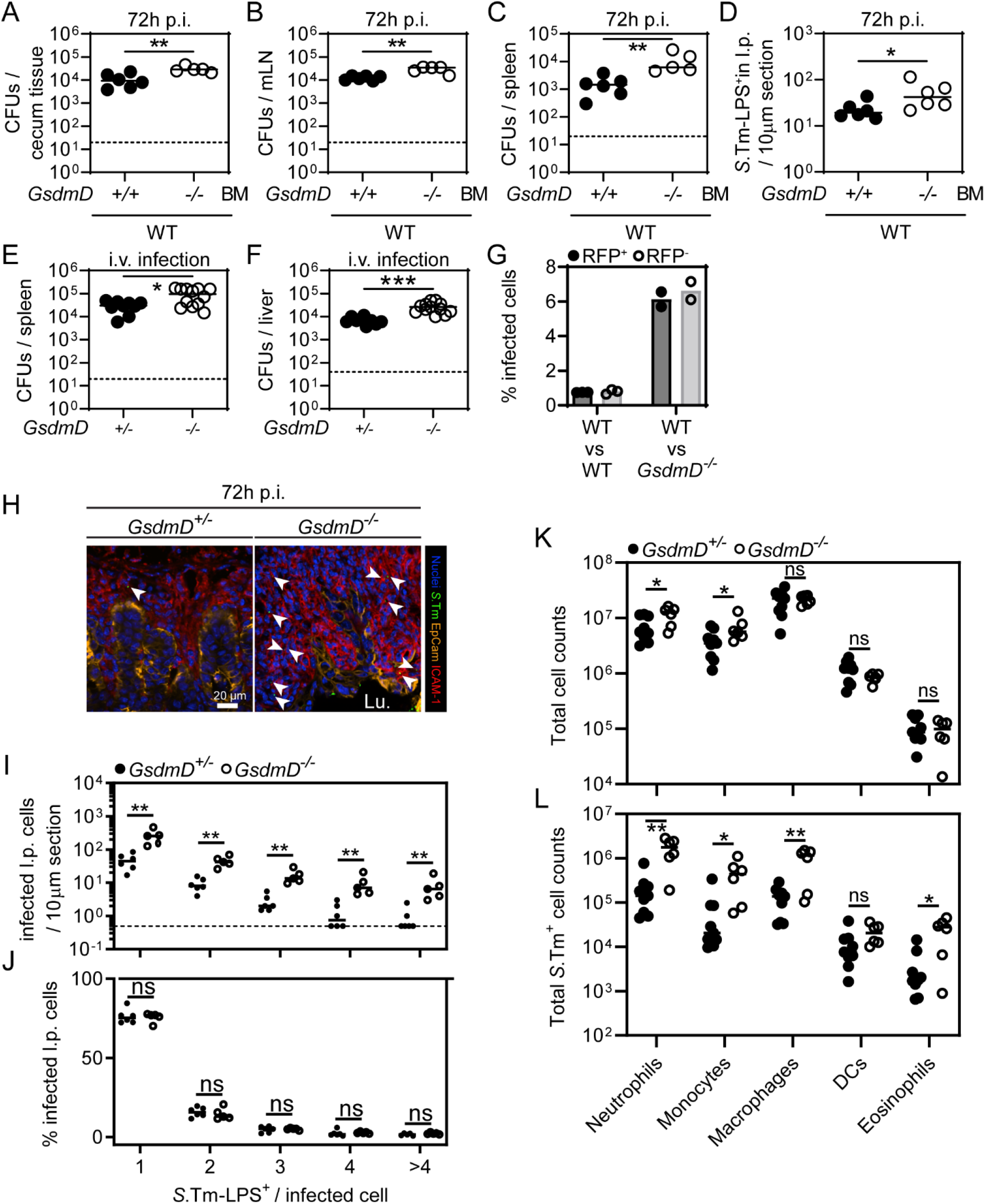
Bone marrow cells in the lamina propria employ GSDMD to restrict *S*.Tm loads. (**A-D**) Bone marrow cells in the lamina propria employ GSDMD to restrict *S*.Tm. Transfer of *GsdmD*^*-/-*^ bone marrow (BM) cells results in elevated *S*.Tm pathogen loads locally and systemically at 72h p.i. *S*.Tm pathogen loads in (**A**) cecum tissue, (**B**) mesenteric lymph nodes, and (**C**) spleen. (**D**) Microscopy-based quantification of *S*.Tm-LPS^+^ cells in the lamina propria. (**E-F**) GSDMD limits *S*.Tm pathogen loads during systemic *S*.Tm infection. *S*.Tm pathogen loads at 24h p.i. of intravenously infected mice in (**E**) spleen, and (**F**)liver. (**G**) *S*.Tm infection of mixed bone marrow chimeras with a 1:1 ratio of either RFP-expressing GSDMD-proficient cells and RFP-non-expressing GSDMD-deficient cells or WT RFP-expressing and - non-expressing cells as a control. Percentage of *S*.Tm-LPS^+^ cells determined by flow cytometry. (**H-J**) GSDMD-deficiency leads to a general increase of infected lamina propria cells. Fluorescence microscopy-based quantification of *S*.Tm-LPS^+^ lamina propria cells at 72h p.i.. (**H**) Representative micrographs of cecum tissue sections, stained for *S*.Tm-LPS. Arrowheads indicate *S*.Tm in lamina propria. Lu. - Lumen. (**I**)Microscopy-based quantification of *S*.Tm-LPS^+^ cells in the lamina propria grouped by the number of *S*.Tm-LPS^+^ per cell. (**J**) Relative percentage of the quantification in K. (**K-L**) Lamina propria cells more frequently harbor *S*.Tm in GSDMD-deficient mice. Flow cytometry analysis of lamina propria cells from 72h infected *GsdmD*^*+/-*^ and *GsdmD*^*-/-*^ littermates. (**K**) Total cell population sizes in lamina propria. (**L**) Total *S*.Tm-LPS^+^ cell numbers in lamina propria. In A-G and I-L each data point represents one mouse. Data is combined from ≥2 independent experiments for each comparison except for G, where only one representative experiment is shown out of 2. Line at median. Dotted line represents detection limit. Mann-Whitney U-test (ns - not significant, *p<0.05, **p<0.01, ***p<0.001).

It is well established that membrane pore-formation by GSDMD induces pyroptosis, which leads to the release of inflammatory mediators including IL1β or IL18, which could act on neighboring cells to prevent *S*.Tm growth. At 48h p.i. we did however not detect elevated systemic *S*.Tm loads in IL18-deficient mice (Fig S4I-J). Furthermore, even in the presence of an IL18-depleting antibody, we still enumerated higher systemic pathogen loads in *GsdmD*^*-/-*^ mice compared to heterozygous littermates, suggesting that GSDMD can limit *S*.Tm independently of IL18 (Fig S4K-L). Similar results with limited numbers of mice were obtained upon depleting IL1β instead (Fig S4M-N), suggesting that neither IL18 nor IL1β mediates the GSDMD-dependent *S*.Tm restriction.

To test if the GSDMD phenotype can be observed on the single cell level, we generated BM chimeras, in which the bone marrow from *GsdmD*^*-/-*^ mice was replaced by a 1:1 mix of RFP expressing WT (*ActRFP*) and non-fluorescent *GsdmD*^*-/-*^ BM cells (Fig S5A). Thereby, we were able to compare GSDMD-deficient and proficient BM-derived cells within the same mouse. As a control we used WT mice and replaced the bone marrow with a 1:1 mix of RFP expressing WT (*ActRFP*) and non-fluorescent WT BM cells (Fig S5A). Notably, flow cytometry analysis of lamina propria cells at 72h p.i. revealed that GSDMD-proficient and -deficient cells in *GsdmD*^*-/-*^ recipients were infected with a similar frequency (Fig 3G, S5B). Hence, in the background of an overall GSDMD-deficient tissue, individual GSDMD-expressing BM-derived cells fail to keep the infection at bay (Fig 3G). This suggests that rather GSDMD having a cell-autonomous impact on *S*.Tm loads, GSDMD-deficiency appears to negatively affect the ability of the mucosal tissue as a whole to control *S*.Tm loads. In line with these observations, we detected both more single and multiple bacterium-containing lamina propria cells in whole-body *GsdmD*^*-/-*^ mice by fluorescence microscopy (Fig 3H-I), while the GSDMD status did not seem to affect intracellular *S*.Tm growth as judged by the relative proportions of lamina propria cells harboring 1, 2, 3, 4 or >4 bacteria, respectively (Fig 3J).

Next, we sought to address if the two highly abundant lamina propria cell types, neutrophils and macrophages, employ GSDMD-dependent restriction mechanism(s). This was of interest, since a role of GSDMD has been shown in these cells (Chen et al., 2018; Ma et al., 2021). We depleted neutrophils with anti-Ly6G or macrophages with anti-CSFR1 antibodies and investigated if *GsdmD*^*-/-*^ mice still featured elevated *S*.Tm organ loads compared to heterozygous littermates. Surprisingly, neither the depletion of neutrophils, nor of macrophages, attenuated the GSDMD-dependent *S*.Tm restriction (Fig S6A-G). Guided by this observation, we explored if GSDMD-deficiency leads to increased *S*.Tm loads in any specific lamina propria cell type. Flow cytometry analysis of lamina propria cells from infected *GsdmD*^*+/-*^ and *GsdmD*^*-/-*^ mice was performed to determine the predominant cell type(s) that harbor *S*.Tm (Fig S6H). Interestingly, while cellular composition was only marginally different between genotypes, multiple cell populations including neutrophils, monocytes and macrophages (which are most frequent and harbor the highest *S*.Tm loads), but also eosinophils (which are less frequent and harbored lower *S*.Tm loads) all featured elevated fractions of *S*.Tm infected cells when comparing GSDMD-deficient mice to their littermate controls (Fig 3K-L). Taken together, GSDMD-deficiency increases *S*.Tm loads across several different types of BM-derived lamina propria cells, particularly in neutrophils, monocytes, macrophages and eosinophils.

### Epithelial and lamina propria GSDMD are both contributing to anti-*S*.Tm defense in the mucosa

The results above demonstrate a restrictive role of GSDMD in lamina propria BM-derived cells. However, it remained unclear if epithelial GSDMD may also contribute to the defense against *S*.Tm, e.g. by cytokine release or by controlling the extrusion of infected IECs. Work in NLRC4-deficient mice had established that these phenotypes are particularly prominent during the first day of the infection of streptomycin pre-treated mice, or the first few hours in enteroid infection models (Sellin et al., 2014; Rauch et al., 2017; Fattinger et al., 2021a; Samperio Ventayol et al., 2021). To tackle this question, we established enteroids from WT and GSDMD-deficient mice, which were infected in-bulk *ex vivo* with a *S*.Tm reporter strain that turns GFP positive upon host cell invasion (*S*.Tm-G^+^, (Hapfelmeier et al., 2005)). *Nlrc4*^*-/-*^ enteroids served as a positive control (Fattinger et al., 2021a; Samperio Ventayol et al., 2021). Interestingly, quantifying *S*.Tm-G^+^ infection foci in >70 enteroids per replicate in our infection experiment (which represents a sufficient sampling size to obtain statistically valid results; Fig S7A) revealed that *GsdmD*^*-/-*^ enteroids harbored significantly more intracellular *S*.Tm than the WT controls. The *S*.Tm counts in the *GsdmD*^*-/-*^ enteroids were, however, still much lower in comparison to NLRC4-deficient enteroids (Fig 4A-B).

**Figure 4.**
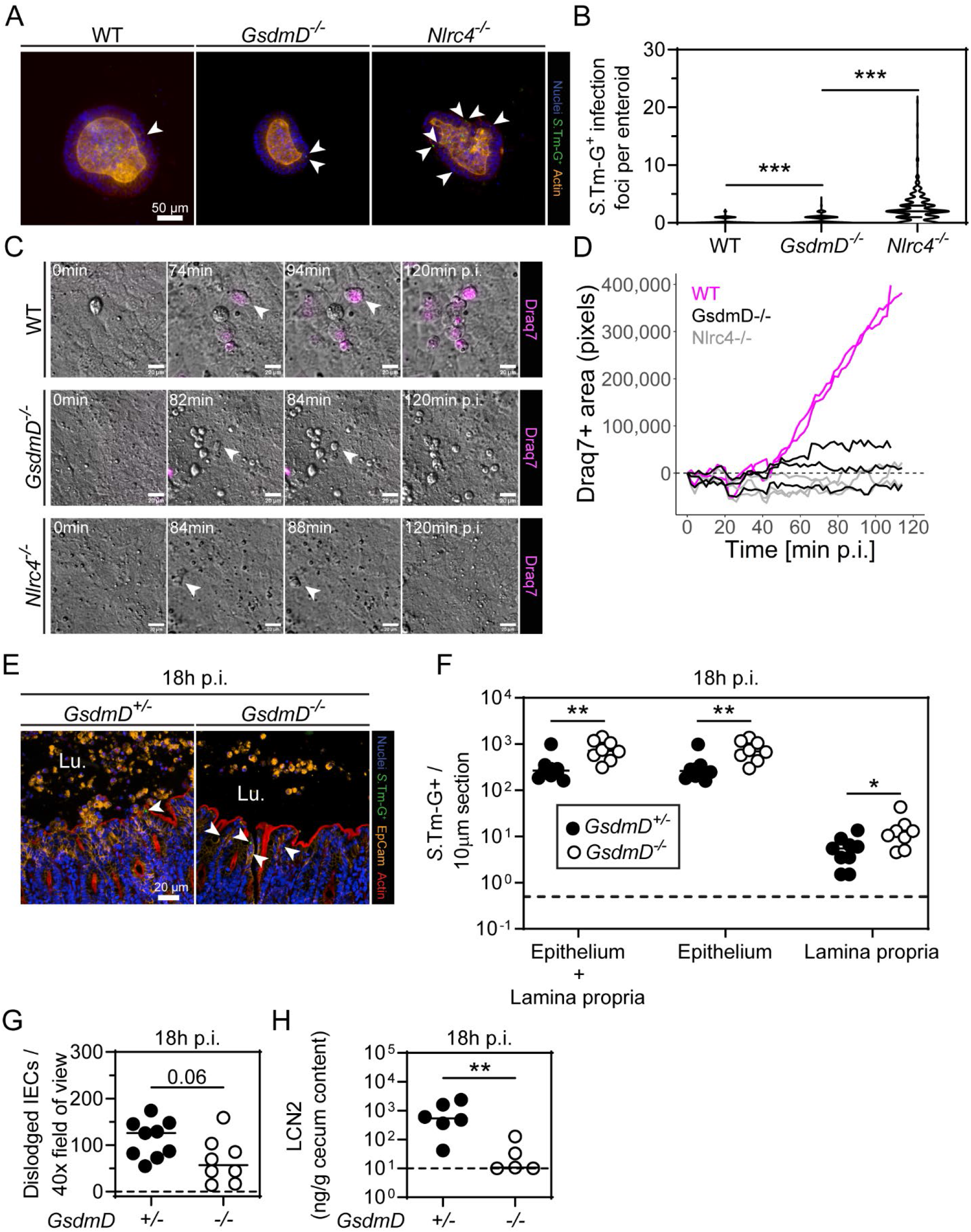
Epithelial GSDMD contributes to the defense against *S*.Tm. (**A-B**) Epithelial GSDMD restricts *S*.Tm loads in epithelium, but not to the same extent as epithelial NAIP/NLRC4. 3D enteroids were infected in bulk with *S*.Tm harboring a *pssaG-GFP* reporter (renders the bacterium GFP-positive upon host cell entry, *S*.Tm-G^+^) for 4h. (**A**) Representative micrographs of infected 3D enteroids. Arrowheads indicate *S*.Tm-G^+^. (**B**) Microscopy-based quantification of *S*.Tm-G^+^ in epithelium of enteroids. (**C-D**) Epithelial GSDMD impacts qualitative features of extruding infected IECs. Enteroid-monolayers were infected with *S*.Tm in the presence of the membrane impermeable dye Draq7 to track membrane lysis. (**C**) Representative micrographs of time-laps microscopy. (**D**) Quantitative analysis of the Draq7 signal from time-laps microscopy. (**E-H**) 18h infections with *S*.Tm harboring a *pssaG-GFP* reporter indicate that epithelial GSDMD restricts *S*.Tm *in vivo* and promotes induction of inflammation. (**E**) Representative micrographs of cecum tissue sections. Arrowheads indicate *S*.Tm-G^+^ in epithelium. Lu. - Lumen. (**F**) Microscopy based quantification of *S*.Tm-G^+^ in mucosal tissue. (**G**) Microscopy based quantification of dislodged IECs. (**H**) Quantification of inflammation by Lipocalin-2 levels of cecum content. In A-D, combined results, or representative results from ≥2 independent experiments. In F-H, each data point represents one mouse. ≥5 mice per group from ≥2 independent experiments for each comparison. Line at median. Dotted line represents detection limit. Mann-Whitney U-test (*p<0.05, **p<0.01, ***p<0.001).

Given that NLRC4 counteracts *S*.Tm infection by expelling infected IECs into the gut lumen (Sellin et al., 2014; Rauch et al., 2017; Fattinger et al., 2021a; Samperio Ventayol et al., 2021), we next sought to address if and how epithelial GSDMD might contribute to this defense mechanism. To this end, we established enteroid-derived monolayers atop loose hydrogels, infected these with *S*.Tm, and followed the IEC extrusion process by differential interference contrast (DIC) and fluorescence live cell microscopy. As expected, NLRC4-deficient monolayers failed to extrude infected IECs (Fig 4C-D). By sharp contrast, in WT and *GsdmD*^*-/-*^ monolayers, we could detect many extruding IECs within 120min of *S*.Tm infection (Fig 4C-D). Importantly, this process was morphologically distinct between WT and GSDMD-deficient monolayers. IECs extruding from the *GsdmD*^*-/-*^ monolayers appeared round and bright in DIC and remained impermeable for the dye Draq7, while extruding WT IECs were as a rule translucent and eventually all became Draq7 positive (Fig 4C-D). Hence, Gasdermin D is not required for extrusion *per se*, but it affects the qualitative features of extruding IECs and, in line with previous work (Rauch et al., 2017; Nozaki et al., 2022), the time point of membrane permeabilization.

To investigate the contribution of epithelial GSDMD *in vivo*, we infected streptomycin pretreated mice for 18h. Based on previous work (Fattinger et al., 2020), it is well established that up to 18h p.i., IECs comprise the predominant infected cell type and determine early disease outcome. Strikingly, while initial luminal colonization was equal between *GsdmD*^*-/-*^ and *GsdmD*^*+/-*^ littermate controls (Fig S7B), we observed increased loads of *S*.Tm-G^+^ in the mucosal tissue and the mLN of *GsdmD*^*-/-*^ mice already at this earlier time point (Fig 4E-F, S7C). This phenotype was attributable to an increased fraction of both infected IECs and lamina propria cells (Fig 4F). In contrast to the profoundly reduced numbers of expelling infected IECs that we had observed in NLRC4-deficient mice (Sellin et al., 2014; Fattinger et al., 2021a), we still observed considerable numbers of dislodged IECs in the GSDMD-deficient mice (Fig 4G). However, GSDMD-deficient mice did feature a trend towards reduced numbers of dislodged IECs and were significantly less inflamed, as judged by lipocalin-2 levels (Fig 4G-H). Of note, despite abundant active Caspase-3 in IECs captured in the extrusion process (Fig S7D), a point mutation rendering GSDMD insensitive to inactivation by Caspase-3, did not alter *S*.Tm tissue loads (Fig S7E-I). Moreover, this gut mucosal phenotype at 18h p.i. was unique to GSDMD and could not be observed in GSDME-deficient mice (Fig S7J-N), not even in a GSDMD-deficient background (Fig S7O-S).

From these observations, we conclude (i) that extrusion of infected IECs can still occur in the absence of GSDMD (and GSDME), (ii) but that epithelial GSDMD dramatically affects the qualitative features of the IECs extrusion process (including the timing of membrane permeabilization) and affects its kinetics. In addition, our combined data demonstrate that both epithelial and lamina propria cell GSDMD contribute to restricting *S*.Tm tissue loads upon oral *S*.Tm infection.

## DISCUSSION

Gasdermins are key executors of multiple pathogen restriction mechanisms. A limited number of *in vivo* studies have shown that Gasdermins C, D, and E can reduce organ loads of diverse pathogens, or are involved in immunopathology (Chen et al., 2021; Demarco et al., 2020; Estfanous et al., 2021; Ma et al., 2021; Tan et al., 2021; Xi et al., 2021; Zhao et al., 2022; Zhang et al., 2022). Nevertheless, to date we still lack a systematic assessment of their potential restrictive role(s) during oral *S*.Tm infection. Our experiments in streptomycin pretreated mice establishes how Gasdermins contribute to *S*.Tm restriction and demonstrate a non-redundant function of GSDMD. Moreover, IECs and lamina propria cells both employ GSDMD to restrict *S*.Tm tissue infection and to switch the gut mucosa as a whole towards an anti-*S*.Tm state.

GSDMD is the best studied Gasdermin to date. GSDMD-deficient mice feature elevated organ pathogen loads during infections with the lung pathogen *Burkholderia cenocepacia* (Estfanous et al., 2021). Moreover, GSDMD also plays an important role in protection against gut pathogens such as *Yersinia pseudotuberculosis* and *Citrobacter rodentium* (Chen et al., 2021; Demarco et al., 2020; Zhang et al., 2022). With regards to *S*.Tm infections, it was reported that GSDMD mediates NETosis in neutrophils upon non-canonical Caspase-11 activation by the attenuated *S*.Tm strain *ΔsifA* (Brinkmann et al., 2004; Chen et al., 2018). *SifA*-deficiency is known to promote egress of *S*.Tm from the *Salmonella* containing vacuole (Beuzón et al., 2000), which should enhance pathogen detection in the host cell’s cytosol by Caspase-11. Interestingly, *GsdmD*^*-/-*^ mice intraperitoneally infected with this mutant strain exhibited elevated pathogen loads in the spleen. This appeared to be dependent on the formation of neutrophil extracellular traps (NETs), since DNase I treatment increased spleen pathogen loads in WT mice, but not in GSDMD-deficient animals. Notably, due to the use of a mutant *S*.Tm strain and the intraperitoneal administration (which bypasses the gut tissue invasion steps of the normal infection process), it remained unclear if this holds true in oral infection with WT *S*.Tm. Here, we demonstrate that GSDMD indeed restricts orally administered WT *S*.Tm in BM-derived lamina propria cells, as well as in systemic organs. However, GSDMD-deficiency in a fraction of BM-derived cells appears enough to increase overall tissue *S*.Tm loads and neutrophils, but also several other immune cells show elevated *S*.Tm numbers in *GsdmD*^-/-^ mice. This suggests that GSDMD acts globally to restrict *S*.Tm in the mucosal tissue. Since neither the separate depletion of neutrophils, macrophages, or the inflammasome-dependent cytokines IL18 or IL1β significantly impacted the GSDMD phenotype, we speculate that multiple mechanisms may explain this restrictive effect of GSDMD. It is likely that GSDMD in macrophages and neutrophils (i) promotes cell death, (ii) accelerates mucosal inflammation, and (iii) traps *S*.Tm in pore-induced intracellular traps (PITs) and NETs, respectively (Brinkmann et al., 2004; Jorgensen et al., 2016), thereby preventing subsequent re-infections into adjecent host cells. These mechanisms are supported by other previous studies *in vivo* (Ma et al., 2021; Chen et al., 2018). Nevertheless, it is plausible that additional mechanism(s) may also contribute to the GSDMD-dependent restriction in the lamina propria.

The data from the infection of 3D enteroids, enteroid-derived monolayers, and early (first 18h) infections in mice suggest that epithelial GSDMD also contributes to restricting *S*.Tm, in particular during the initial phase of the infection. Importantly, this restriction is much weaker compared to that conferred by epithelial NAIP/NLRC4. This can be explained by the prominent role for NAIP/NLRC4 in driving extrusion of infected IECs, a process which can still be executed also in the absence of GSDMD. However, in GSDMD-deficient enteroid-monolayers we found that the qualitative features of extruding cells appear remarkably different. In line with a very recent publication, we show that epithelial GSDMD impacts the time point of membrane lysis during the extrusion process, which also influences how efficiently, and with what kinetics, infected IECs can be removed from the epithelium (Nozaki et al., 2022). Furthermore, a delay of cell membrane permeabilization towards much later time points (that is at or after the end of the IEC extrusion process) should also impact the levels of pro-inflammatory mediators, such as IL1β, IL18, and inflammatory lipids, that can reach the lamina propria to induce inflammation. In fact, at 18h p.i. we do observe a delayed onset of inflammation in GSDMD-deficient mice *in vivo*, which may be a result of reduced exposure of the lamina propria to IEC-derived pro-inflammatory cytokines. Notably, another recent study showed that goblet cells rely on GSDMD to secrete mucus on top of the epithelium, and that this may shield against microbes (Zhang et al., 2022). However, in the unperturbed gut, we did not observe any change in the steady-state level of inflammatory marker genes in the cecum tissue of *GsdmD*^-/-^ mice. This is in line with that the mucus covers only the bottom of the crypts of the murine cecum epithelium (which is the main site of *S*.Tm attack in the guts of streptomycin pretreated mice), while the epithelial cells at the tip of the crypts are lacking such mucus cover, as observed in earlier work on wild type mice (Furter et al., 2019). In either case, our combined results highlight how epithelial and immune cell GSDMD complement each other in the defense of the mucosal tissue against *S*.Tm invasion.

Recent work in mice has demonstrated that other Gasdermins, e.g. GSDMC and GSDME, also take part in the defense against pathogens and in inflammation (Chen et al., 2021; Tan et al., 2021; Xi et al., 2021). In particular, GSDME was shown to induce Caspase-8 driven pyroptosis in neutrophils, which helps to control systemic *Yersinia* infections (Chen et al., 2021). Additionally, GSDME in epithelial cells promotes inflammation during chemically induced colitis (Tan et al., 2021). Therefore, it is somewhat surprising that we do not detect any protective effect for GSDME during *S*.Tm infection. GSDME seems to neither influence IEC extrusion efficiency, nor to restrict *S*.Tm pathogen loads in the lamina propria, or at systemic sites, not even in a GSDMD-deficient background. Furthermore, GSDMC2 and GSDMC3, which were shown to be highly expressed in epithelial cells of worm-infected mice (Xi et al., 2021), along with GSDMC1 and GSDMC4 also fail to impact *S*.Tm infection. The same holds true for GSMDA1-3. Why only GSDMD plays an important protective role during *S*.Tm infection is not fully clear. A reason could be that innate immunity against *S*.Tm is dominated by inflammasome signaling or that there is redundancy for GSDMA1-3, C1-4 and E and that only combined deficiency might reveal a phenotype. Also, we cannot rule out that *S*.Tm may express yet unidentified virulence factors blocking the action of some Gasdermins. As GSDMD is the main Gasdermin cleaved by Caspase-1, and as Caspase-1 is an important factor contributing to NAIP/NLRC4-mediated protection of the intestinal mucosa against *S*.Tm, it makes sense that this specific Gasdermin mediates restriction.

In summary, we have assessed the role of Gasdermins in the defense against acute oral *S*.Tm infection. Our work demonstrates that out of all Gasdermins, only GSDMD exerts a significant restrictive function. Both the IEC and lamina propria defense systems notably rely on GSDMD, which contributes to multiple restrictive mechanisms across these two compartments. It remains to be shown, if this is specific for the pathogen investigated here, or of broader relevance for the mucosal defense against other invasive enteropathogenic bacteria.

## METHODS

### Bacterial strains, plasmids and culture conditions

All infection experiments were done with *Salmonella* Typhimurium (*S*.Tm) SL1344 (SB300, SmR) if not otherwise specified. Where indicated, *S*.Tm reporter strain harboring the plasmid pM975 (*passaG*-GFPmut2) was used (Hapfelmeier et al., 2005). *S*.Tm was cultured overnight in LB/0.3M NaCl (Sigma Aldrich) with appropriate antibiotics (ca. 12h) before sub-culturing in 1:20 dilution for 4h in the same media without antibiotics. For mouse infections, *S*.Tm were washed once and reconstituted with PBS (BioConcepts) before oral gavage. For 3D enteroid infections, *S*.Tm were washed with PBS and reconstituted in DMEM/F-12 (STEMCELL) supplemented with 3% FCS (Thermo Fisher). For 2D enteroid-derived monolayer infections, *S*.Tm were reconstituted in DMEM/F-12 (Gibco) and diluted in complete mouse Intesticult (STEMCELL) without antibiotics to the desired concentration.

### Mouse infections

All mice used were specific pathogen free (SPF) and were maintained in individually ventilated cages of the ETH Zürich mouse facility (EPIC and RCHCI). WT mice were C57BL/6 originally from Charles River (Sulzfeld, Germany). All genetic modified mice were of C57BL/6 background. Specifically, the following mouse lines were used: *Nlrc4*^*-/-*^ (B6.C2-Nlrc4tm1Vmd, (Mariathasan et al., 2004)), *GsdmD*^*-/-*^ (Shi et al., 2015), *GsdmD*_fsX^*-/-*^ (C57BL/6J-*Gsdmd*^*em1Broz*^, (Chen et al., 2019)), *GsdmD*^*D88A/D88A*^ (C57BL/6J-*Gsdmd*^*em2Broz*^, (Chen et al., 2019)), *GsdmE*^*-/-*^ (C57BL/6J-*Gsdme*^*em1Broz*^, (Chen et al., 2019)), *GsdmD*^*-/-*^*GsdmE*^*-/-*^ (C57BL/6J-*Gsdmd*^*em1Broz*^*Gsdme*^*em1Broz*^, (Chen et al., 2021)), *GsdmA*^*-/-*^ (C57BL/6J-*Gsdma1-3*^*em1Broz*^, this study), *GsdmC*^*-/-*^ (C57BL/6J-*Gsdmc1-4*^*em1Broz*^, this study), *ActRFP* (B6.Cg-Tg[CAG-DsRed*MST]1Nagy/J, (Vintersten et al., 2004)), *IL18*^*-/-*^ (B6.129P2-Il18tm1Aki, (Takeda et al., 1998)). Genotyping of mice was done by PCR or sequencing. Heterozygous littermates were used as control animals. 8-14 weeks old mice were infected according to the streptomycin mouse model (Barthel et al., 2003). Briefly, mice were orally pretreated with 25mg streptomycin sulfate (Sm, Applichem) one day before infection with ∼5×10^7^ CFU *S*.Tm by oral gavage. Mice were monitored daily, and organs were harvest at the indicted time points. Organs were homogenized in PBS containing 0.5% tergitol, 0.5% BSA using a tissue lyser (Qiagen) and plated on MacConkey agar (Oxoid) with Sm. Cecum tissue was first washed in PBS, incubated for 30-60min in PBS/400μg/ml gentamycin and washed extensively (6x) in PBS before plating. To generate bone marrow chimeras, mice were gamma-irradiated (1000 Rad, 14min) and 5×10^6^ bone marrow cells from the respective mouse line was transferred via tail vein. Mice received Borgal (Vererinaria AG) in the drinking water for 3 weeks and kept at least for 8 weeks before infection. For intravenous infections, 10^4^ *S*.Tm in 100ul PBS from an overnight or 4h subculture were injected in the tail vein. For IL18 or IL1β depletion experiments, 200ug/mouse anti-IL18 (BioXCell, YIGIF74-1G7) or anti-IL1β (BioXCell, B122), respectively, were injected intraperitoneal on the day of pretreatment and infection. For neutrophil depletion experiments, 500μg/mouse anti-Ly6G (BioXCell, 1A8) was injected intraperitoneal daily starting at pretreatment. For macrophage depletion, 1000μg/mouse anti-CSFR1 (BioXCell, AFS98) was injected intraperitoneal 4 days prior infection and 300μg/mouse every following day until harvesting. All animal experiments were approved by the Kantonales Veterinäramt Zürich (licences 193/2016 and 158/2019).

### Murine 3D intestinal epithelial enteroids and infections

Murine jejunal epithelial enteroids were established and maintained as previously described (Fattinger et al., 2021a). Briefly, 2mm pieces of the mouse jejunum were washed in ice-cold PBS, incubated in Gentle cell dissociation reagent (STEMCELL) while rocking (20rpm, 15min, RT), and transferred to PBS/0.1%BSA (Chemie Brunschwig AG) to extract intestinal crypts by mechanical shearing. Extracted crypts were filtered through a 70μm cell strainer, washed, and embedded in 50μl Matrigel (Chemie Brunschwig AG) domes. The enteroids were maintained in complete mouse Intesticult medium (STEMCELL) supplemented with PenStrep (Gibco) (37°C, 5% CO_2_). The culture medium was exchanged every 2-4 days. Every 5-7 days, the cultures were split by mechanical shearing in RT Gentle dissociation reagent and the enteroids were re-embedded in 50ul Matrigel domes (splitting ratio 1:4 to 1:6). Stable enteroid cultures were cryopreserved and thawed for experimentation. *S*.Tm bulk infections were performed after at least 2 weeks of culture maintenance. To this end, domes containing ∼100 enteroids were dissolved in ice-cold DMEM/F-12/3%FCS by pipetting carefully up and down. The enteroids were pelleted by centrifugation (300g, 5min, 4°C), and re-suspended in pre-warmed DMEM/F-12/3%FCS without antibiotics. *S*.Tm harboring a *pssaG-GFP* reporter was used to infect enteroids at an estimated MOI of 100 for 40min (37°C, 5% CO_2_) (assumption of ∼1000 epithelial cells per enteroid (Fattinger et al., 2021a). After infection, RT DMEM/F-12/3% FCS containing 100μg/ml gentamycin (AppliChem) was added for 15min (37°C, 5% CO_2_) to kill extracellular bacteria. The enteroids were pelleted (300g, 5min, 4°C), resuspended in complete Intesticult supplemented with 25μg/ml gentamycin, and seeded in 25μl Matrigel domes in prewarmed 8-well glass chamber slides (Thermo Scientific). Domes were solidified for 10min (37°C, 5% CO_2_), RT complete IntestiCult containing 25μg/ml was added, and enteroids were incubated (37°C, 5% CO_2_) until fixation with 4% paraformaldehyde (PFA; Sigma Aldrich) at 4h p.i. After fixation, samples were washed three times with PBS, permeabilized with PBS/0.5% Tx-100 (Sigma Aldrich) for ≥10min, blocked with PBS/10% Normal Goat Serum (NGS) for ≥30min, and incubated for ≥ 40min with TRITC-conjugated Phalloidin (Fluoprobes) and DAPI (Sigma Aldrich). Stained enteroids were extensively washed with PBS and ddH_2_O, the chambers were carefully removed from the glass slides, and the samples were covered with a glass slip using one drop of Mowiol (VWR International AG) per dome.

### Establishment and infection of 2D murine enteroid-derived monolayers

2D murine enteroid-derived monolayers were established as previously described (Samperio Ventayol et al., 2021). In brief, enteroids were split as described above and cultured in freshly prepared CV medium, i.e. complete mouse Intesticult supplemented with 3μM CHIR99021 (Cayman Chemicals) and 1mM valproic acid (Cayman Chemicals) for one week. The medium was exchanged for fresh CV medium every 2-3 days. Glass-bottom 8-well chamber slides (Cellvis) were pre-coated with 75μg/mL Poly-L-Lysine (Sigma Aldrich) at RT overnight and washed 3 times with PBS (Gibco). Chamber slides were then dried for 2h before a 1mg/mL collagen (Corning) solution in collagen neutralization buffer (20mM HEPES/53mM sodium bicarbonate/sodium hydroxide equimolar to acetic acid from the collagen stock) was added to the wells and the hydrogels were left to solidify for 1h at 37°C as previously described (Hinman et al., 2021). CV-pre-treated enteroids were dissociated by mechanical shearing and incubation in Gentle cell dissociation reagent (STEMCELL) for 10min, nutating at RT. After washing in ice-cold DMEM/F12 (Gibco)/0.25% BSA (Gibco), enteroids were reconstituted in ice-cold DMEM/F12/0.25% BSA and passed through a pre-wetted G25 needle approximately 10 times for mechanical dissociation. Finally, the cells suspension was reconstituted in RT CV medium/10μM Y-27632 (Sigma Aldrich) and 150’000 cells/cm^2^ were added to the prepared collagen I hydrogels. Monolayers were maintained at 37°C, 5% CO_2_, and 24h after establishment, they were washed once in pre-warmed DMEM/F12 and the medium was exchanged for complete mouse Intesticult without Y-27632. Thereafter, the medium was exchanged for fresh complete mouse Intesticult every 1-2 days. Monolayer infections were performed 72-96h post establishment. Prior to infection, the monolayers were washed once with pre-warmed DMEM/F12 and complete Intesticult without antibiotics containing 1.5μM Draq7 was added to each well. After placing the chamber slide in the pre-warmed microscope chamber (37°C, 5% CO_2_), the prepared *S*.Tm inoculum was added at an MOI of 1-2 and imaging was started immediately.

### Time-lapse imaging of 2D murine enteroid-derived monolayers

Time-lapse imaging of 2D murine enteroid-derived monolayers was performed on a custom-built microscope based on an Eclipse Ti2 body (Nikon) with a 60x, 0.7 numerical aperture Plan Apo Lambda air objective (Nikon) and a back-lit sCMOS camera (pixel size 11μm, Prime 95B; Photometrics). Samples were maintained at 37°C, 5% CO_2_ in a moisturized chamber during imaging. Bright-field imaging was performed using differential interference contrast (DIC), and fluorescence was acquired with an excitation light engine Spectra-X (Lumencor) and emission collection through a quadruple bandpass filter (89402; Chroma). Infected monolayers were imaged at 2min intervals for a total of 120min. To quantify IEC permeabilization in response to infection, images were thresholded in Fiji (a version of ImageJ; (Schindelin et al., 2012) using the same threshold value for all time-lapse movies from the same experiment, and the area above threshold was enumerated.

### Immunofluorescence staining, wide field- and confocal microscopy

Upon harvesting, mouse cecum tissue was fixed in 4% PFA, saturated in 20% sucrose, and submerged in Optimal Cutting Temperature compound (OCT, Tissue-Tek) before flash freezing in liquid nitrogen. Samples were kept at -80°C until further analysis. Ceca were cut in 10-20μm thick cross-sections and mounted on glass slides (Superfrost++, Thermo Scientific). Air-dried sections were rehydrated with PBS, permeabilized with PBS/0.5% Tx-100 (Sigma Aldrich) and incubated with PBS/10% Normal Goat Serum (NGS; Reactolab SA) before fluorescence staining. For fluorescence staining the following primary/secondary antibodies and dyes diluted in PBS/10%NGS were used: α-EpCam/CD326 (clone G8.8, Biolegend), α-cleaved Caspase 3 (#9661, Cell Signaling Technology), α-*S*.Tm LPS (O-antigen group B factor 4-5, Difco), α-ICAM-1/CD54 (clone 3E2, BD Biosciences), α-rabbit-AlexaFluor488 (Abcam Biochemicals), α-rabbit-Cy3 (Bethyl Laboratories), α-rat-FITC (Jackson), α-rat-Cy3 (Jackson), α-rat-Cy5 (Jackson), α-hamster-Cy5 (Jackson), CruzFluor488-conjugated Phalloidin (Santa Cruz Biotechnology), TRITC-conjugated Phalloidin (Fluoprobes), AlexaFluor647-conjugated Phalloidin (Molecular Probes) and DAPI (Sigma Aldrich). Mowiol (VWR International AG) was used to cover the stained sections with a coverslip. Microscopy was performed using a Zeiss Axiovert 200m microscope with 10-100x objectives, a spinning disc confocal lased unit (Visitron), and an Evolve 512 EMCCD camera (Photometrics). Images were processed or analyzed with Visiview (Visitron) and/or ImageJ. Microscopy quantification was done manually and blindly on at least 2 sections per mouse as previously described (Fattinger et al., 2021a).

### Flow-cytometric analysis of lamina propria cells

Cecum lamina propria cells were isolated and stained as previously described (Hausmann et al., 2020). For cell surface staining, cells were incubated in 1μg/sample Mouse BD Fc Block (BD Biosciences) in 75 μl 10% Brilliant stain buffer (BD Biosciences)/FACS buffer for 5 min at 4°C prior to adding 25ul of antibody mix in 10% Brilliant stain buffer/FACS buffer. The following antibodies and dyes were used: CD45-PerCP (Biolegend; 30-F11; 1:100), CD45-BUV563 (BD Biosciences; 30-F11; 1:100), MHCII-BV421 (Biolegend; M5/114.15.2; 1:100), CD11c-PE/Cy7 (Biolegend; N418; 1:200), CD3-BV711 (Biolegend; 145-2C11; 1:200), NK1.1-BV711 (Biolegend; PK136; 1:200), B220-BV711 (Biolegend; RA3-6B2; 1:200), Siglec-F-APC/Cy7 (BD Biosciences; E50-2440; 1:200), Ly-6G-BV650 (Biolegend; 1A8; 1:100), Ly6C-AF700 (Biolegend; HK1.4; 1:200), CD64-PE/Dazzle (Biolegend; X54-5/7.1; 1:100), LIVE/DEAD Fixable Aqua Dead Cell Stain (Life Technologies; 1:1,000). For the intracellular *S*.Tm-LPS staining using α-*S*.Tm LPS (O-antigen group B factor 4-5, Difco) and α-rabbit-AlexaFluor647 (Abcam Biochemicals), cells were incubated with IC fixation buffer (eBioscience, 00-8222) and permeabilization buffer (eBioscience, 00-8333). To set the gates for *S*.Tm-LPS^+^ cells, lamina propria cells were only stained with secondary antibody. Samples were measured on a LSRII (BD Biosciences) or LSR Fortessa (BD Biosciences), and data were analyzed with FlowJo V10 (TreeStar).

### Histology

For histology analysis, cecum tissue embedded in OCT was snap frozen in liquid nitrogen, cut in 5μm sections, air-dried and stained with hematoxylin and eosin. Histology score was determined blindly as described previously (Barthel et al., 2003).

### Lipocalin-2 and TNF ELISA

Feces or cecum content was used for Lipocalin-2 ELISA and ca. a 5mm piece of extensively washed cecum tissue for TNF ELISA. Cecum tissue sample was homogenized in PBS/0.5%Tergitol/0.5%BSA (Sigma Aldrich, Chemie Brunschwig AG) supplemented with protease inhibitor cocktail (Roche). Lipocalin-2 (R&D Systems) and high sensitivity TNF (Invitrogen) ELISA was performed according to the manufacturer’s protocols.

### RT-qPCR

Cecum tissue samples were snap frozen in RNAlater (Invitrogen) and kept at -80°C. RNeasy Mini Kit (Qiagen) was used to isolate RNA and RT^2^ HT First Strand cDNA Kit (Qiagen) to reverse transcribe to cDNA. A QuantStudio 7 Flex StepOne Plus Cycler was used to perform qPCR analysis with FastStart Universal SYBR Green Master reagents (Roche). Only validated primers from Qiagen were used.

### Statistical analysis

Mann-Whitney U-test was used to assess statistical significance where applicable as indicated in the figure legends.

## ACKNOWLEDGEMENTS

We thank members of the Hardt, Sellin, and Broz laboratories for helpful discussions. We acknowledge Prachi Shukla for her help during her rotation in the Hardt lab, Dmitri Kotov for his help analyzing the FACS data, the staff of the ETH Zürich mouse facility EPIC/RCHCI (especially Manuela Graf, Katharina Holzinger, Dennis Mollenhauer & Dominik Bacovcin), the UNIL animal facility and the Center for transgenic models (CTM-Basel). This work was financed by grants from the Swiss National Science Foundation (310030_192567) and the Novartis Research Foundation (#21B079) to WDH, and ERC (ERC-2017-CoG 770988, InflamCellDeath) and SNSF (310030_198005) to P.B. EG was funded by a grant from the Monique Dornonville de la Cour Foundation to WDH. MES acknowledges financial support from the Swedish Research Council (2018-02223), and the Swedish Foundation for Strategic Research (ICA16-0031; FFL18-0165).

## AUTHOR CONTRIBUTIONS

Conceptualization: SAF, PB, MES, WDH. Methodology: SAF, LM, PG, EG, SK. Investigation: SAF, LM, PG, UE, SG, EG, SK, BD. Technical assistance: MF, MB. Developed and provided reagents: PP, FS, PB. Writing - Original Draft: SAF. Writing - Review & Editing: SAF, LM, PG, MES, WDH. Visualization: SAF, LM, PG, SK. Funding acquisition: PB, MES, WDH. All authors read, commented and approved this manuscript.

## SUPPLEMENTARY FIGURE

**Figure S1.**
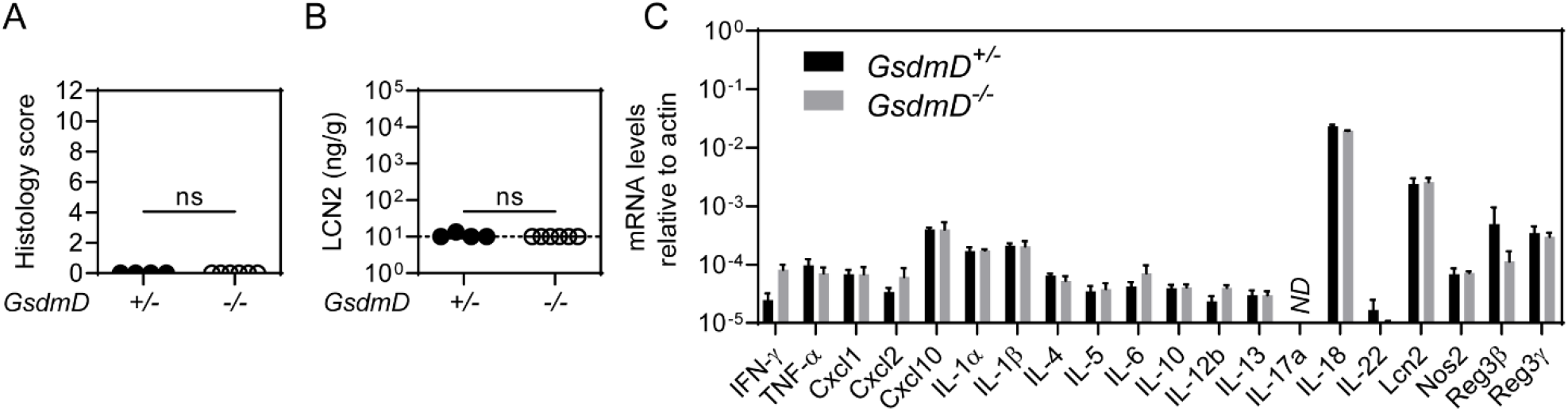
(Steady state analysis of inflammation) (**A-B**) No detectable inflammatory pre-condition in the gut mucosa of GSDMD-deficient mice. (**A**) Microscopy-based quantification of histology score from H&E-stained cecum tissue section. (**B**) Lipocalin-2 levels of feces. In A, B, each data point represents one mouse. Line at median. Dotted line represents detection limit. (**C**) mRNA levels of pro-inflammatory cytokines and anti-microbial peptides. Means with SD are indicated. For all panels ≥4 mice per group. Mann-Whitney U-test (ns - not significant).

**Figure S2.**
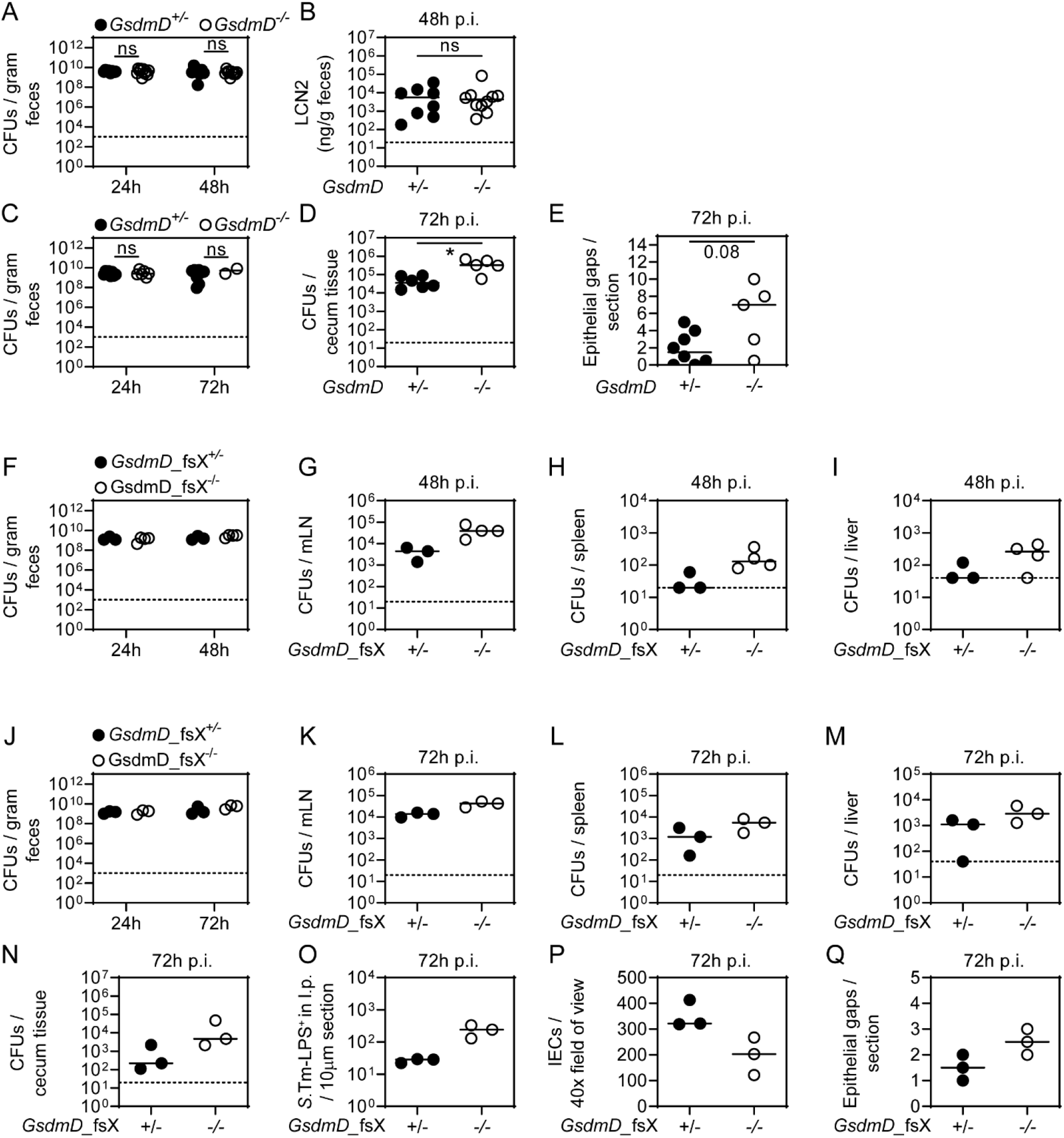
(Supplementary figure to Fig 1) (**A-B**) 48h infection of *GsdmD*^*+/-*^ and *GsdmD*^*-/-*^ littermates. (**A**) *S*.Tm pathogen loads in feces over time. (**B**) Lipocalin-2 levels of feces at 48h p.i. (**C-D**) 72h infection of *GsdmD*^*+/-*^ and *GsdmD*^*-/-*^ littermates. (**C**) *S*.Tm CFU pathogen loads in feces over time. (**D**) *S*.Tm CFU pathogen loads in cecum tissue at 72h p.i. (**E**) Microscopy-based quantification of epithelial gaps in the cecum tissue at 72h p.i. (**F-Q**) Independent GSDMD-deficient mouse line (cause by genetic frameshift, *GsdmD_fsX*) exhibits same phenotype. (**F-I**) 48h infection of *GsdmD_*fsX^*+/-*^ and *GsdmD_*fsX^*-/-*^ littermates. *S*.Tm pathogen loads in (**F**) feces over time and at 48h p.i. in (**G**) mesenteric lymph nodes, (**H**) spleen, and (**I**) liver. (**J-Q**) 72h infection of *GsdmD_*fsX^*+/-*^ and *GsdmD*_fsX^-/-^ littermates. *S*.Tm pathogen loads in (**J**) feces over time and at 72h p.i. in (**K**) mesenteric lymph nodes, (**L**) spleen, (**M**) liver, and (**N**) cecum tissue. Microscopy-based quantification at 72h p.i. of (**O**) *S*.Tm-LPS^+^ cells in the lamina propria, (**P**) IECs per field of view, and (**Q**) epithelial gaps per section. In all panels each data point represents one mouse. ≥3 mice per group. Line at median. Dotted line represents detection limit. Mann-Whitney U-test (ns - not significant, *p<0.05).

**Figure S3.**
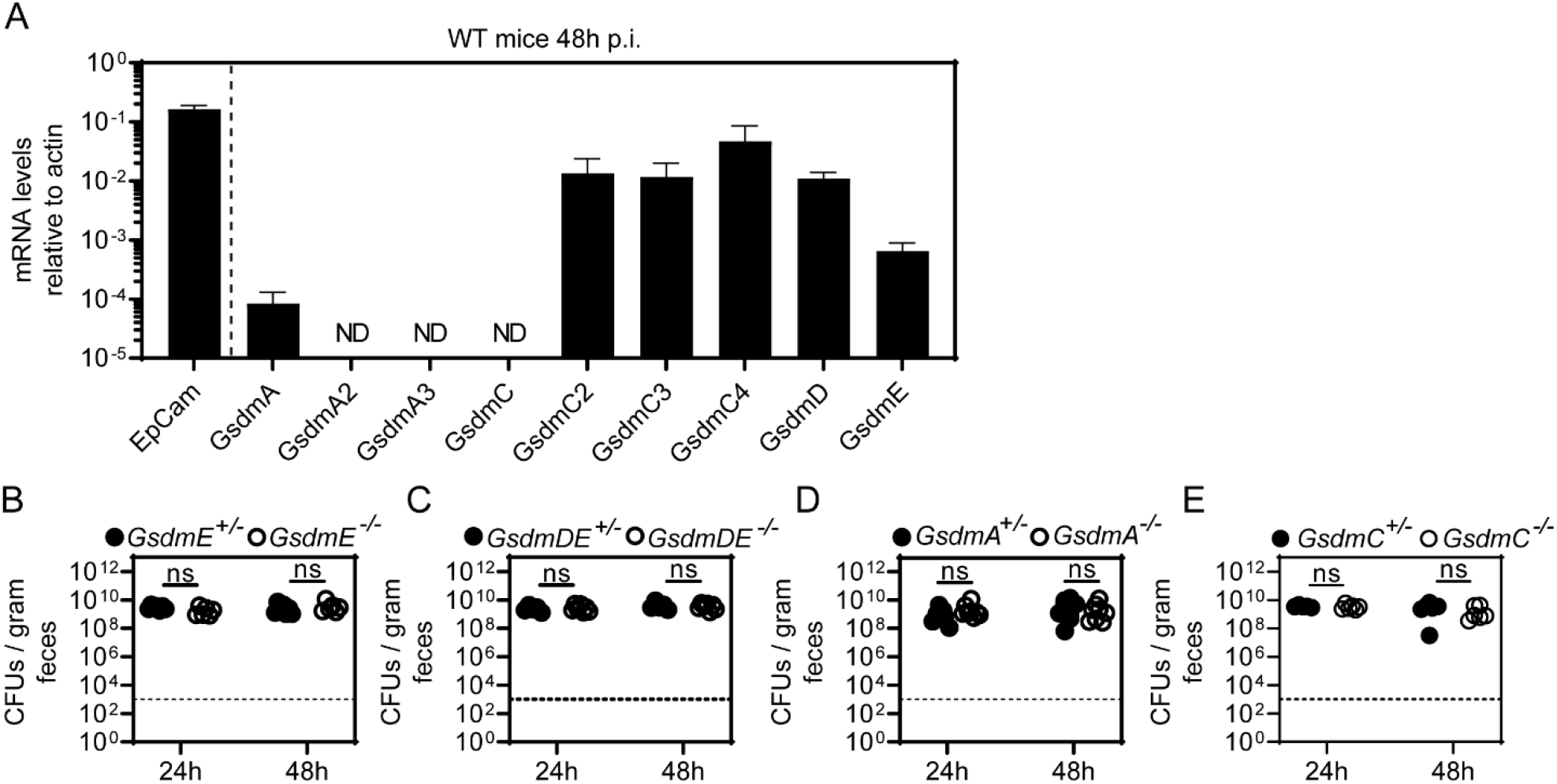
(Supplementary figure to Fig 2) (**A**)Relative expression levels of individual Gasdermins in the cecum tissue at 48h p.i.. (**B-E**) *S*.Tm pathogen loads in feces over time of 48h infections with (**A**) GSDME-deficient, (**B**) GSDMD/E-double-deficient, (**C**) GSDMA-deficient and (**D**) GSDMC-deficient mice. Each data point represents one mouse. ≥5 mice per group from ≥2 independent experiments for each comparison. Line at median. Dotted line represents detection limit. Mann-Whitney U-test (ns - not significant).

**Figure S4.**
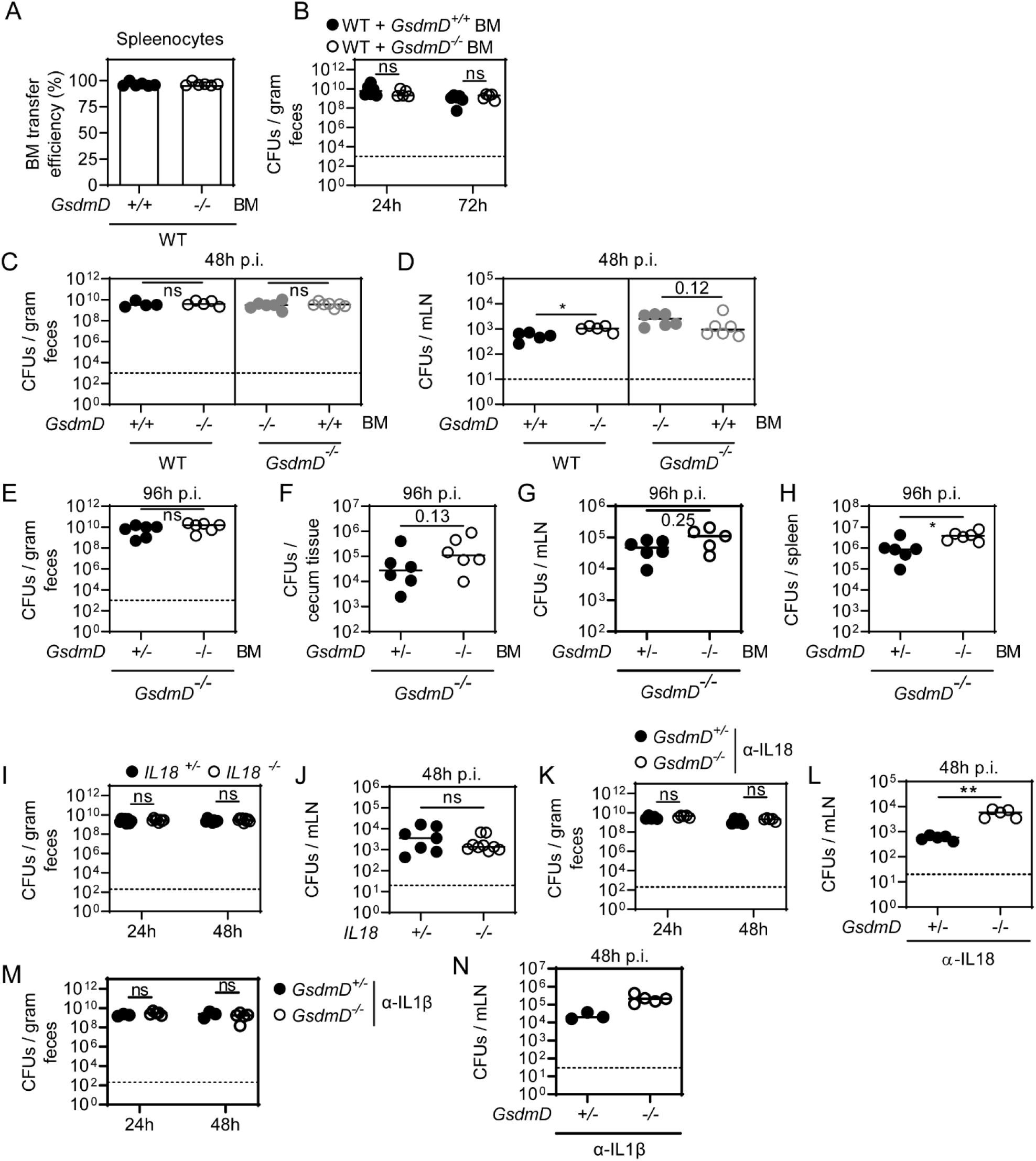
(Supplementary figure to Fig 3) (**A-B**) 72h infection of bone marrow (BM) chimeras. (**A**) Bone marrow transfer efficiency for spleenocytes. (**B**)*S*.Tm pathogen loads in feces over time. (**C-H**) *GsdmD*^*-/-*^ bone marrow (BM) results in elevated *S*.Tm pathogen loads locally and systemically at 48 and 96h p.i. (**C-D**) 48h infection of bone marrow chimeras with *S*.Tm harboring a *pssaG-GFP* reporter. *S*.Tm pathogen loads in (**C**) feces, and (**D**) mesenteric lymph nodes. (**E-H**) 96h infection of bone marrow chimeras with *S*.Tm. *S*.Tm pathogen loads in (**E**) feces, (**F**) cecum tissue, (**G**) mesenteric lymph nodes, and (**H**) spleen. (**I-J**) At 48h p.i., IL18-deficient mice exhibit similar *S*.Tm pathogen loads in (**I**) feces, (**J**) mesenteric lymph nodes. (**K-L**) IL18 is dispensable for GSDMD-dependent *S*.Tm restriction. *S*.Tm pathogen loads at 48h p.i. (**K**) in feces (**L**) in mesenteric lymph nodes of GsdmD^+/-^ and GsdmD^-/-^ littermates in the presence of a neutralizing IL18 antibody. Note, this infection was performed with *S*.Tm harboring a *pssaG-GFP* reporter which explains the overall ca. 10x lower pathogen loads. (**M-N**) IL1β is dispensable for GSDMD-dependent *S*.Tm restriction. *S*.Tm pathogen loads at 48h p.i. (**M**) in feces (**N**) in mesenteric lymph nodes of GsdmD^+/-^ and GsdmD^-/-^ littermates in the presence of a neutralizing ILβ antibody. Each data point represents one mouse. Line at median. Dotted line represents detection limit. Mann-Whitney U-test (ns - not significant, *p<0.05, **p<0.01, ***p<0.001).

**Figure S5.**
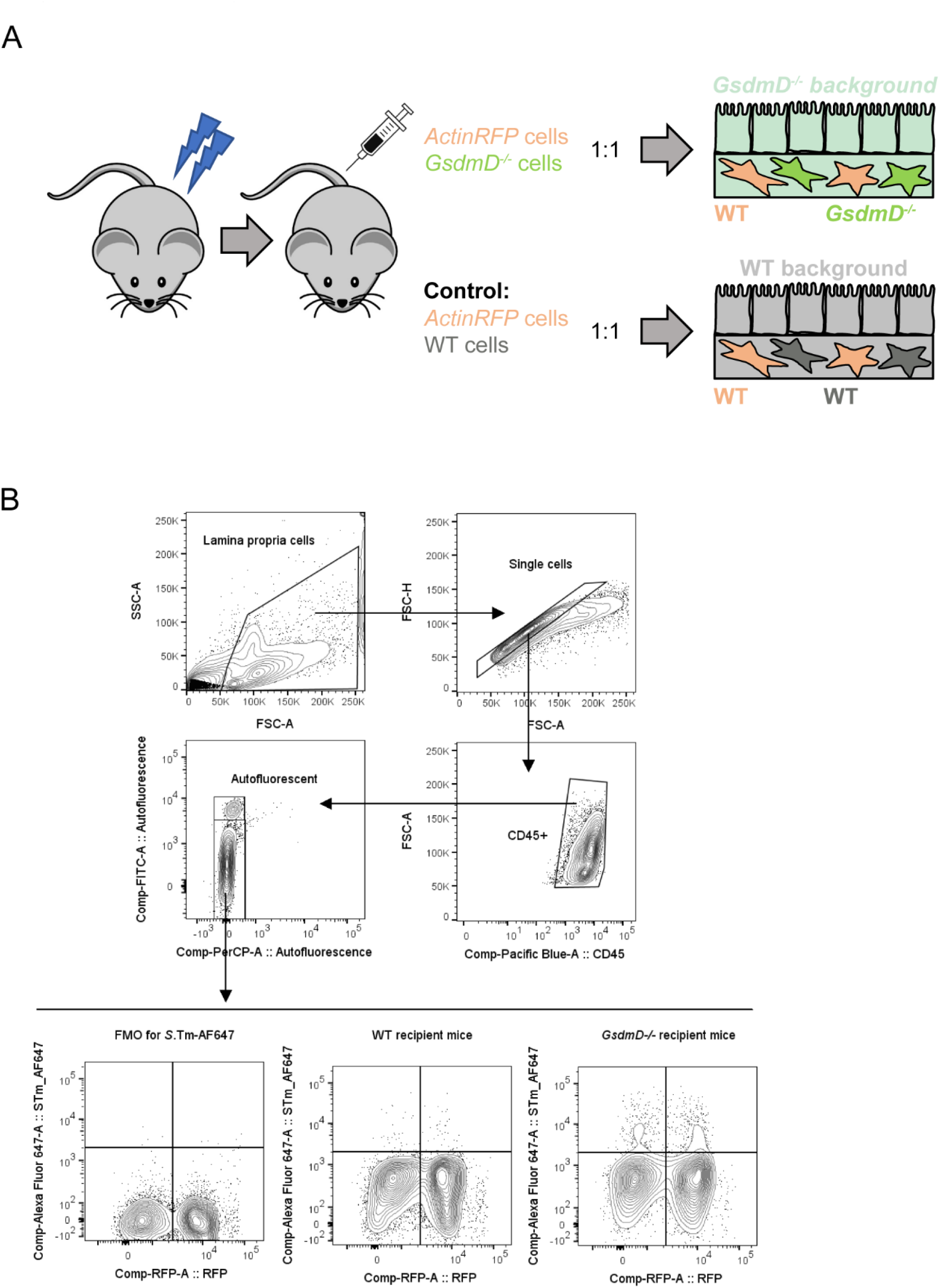
(Supplementary figure to Fig 3) (**A**) Graphical illustration of experimental setup. (**B**) Flow cytometry gating strategy for Fig 3G.

**Figure S6.**
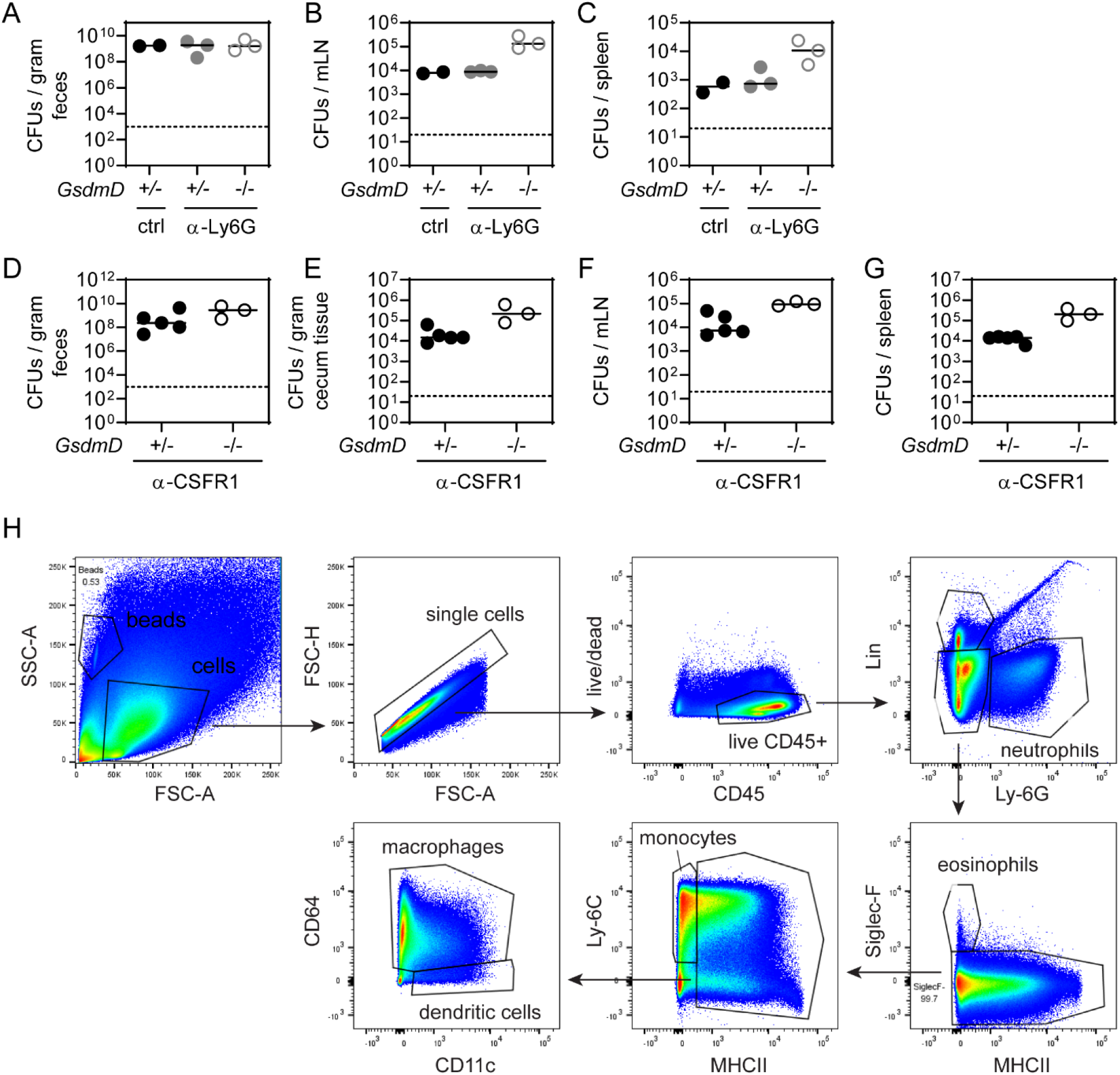
(Supplementary figure to Fig 3) (**A-G**) Depletion of neutrophils nor macrophages eliminates GSDMD-dependent phenotype. (**A-C**) Neutrophil depletion by anti-Ly6G antibody in a 48h *S*.Tm infection of *GsdmD*^*+/-*^ and *GsdmD*^*-/-*^ littermates. Control mice were injected with PBS. *S*.Tm pathogen loads in (**A**) feces, (**B**) mesenteric lymph nodes, and (**C**) spleen. (**D-G**) Macrophage depletion by anti-CSFR1 antibody in a 72h *S*.Tm infection of *GsdmD*^*+/-*^ and *GsdmD*^*-/-*^ littermates. *S*.Tm pathogen loads in (**D**) feces, (**E**) cecum tissue, (**F**) mesenteric lymph nodes, and (**G**) spleen. In all panels each data point represents one mouse. Line at median. Dotted line represents detection limit. (**H**) Gating strategy of flow cytometry analysis to determine lamina propria cell types in figure 3K-L.

**Figure S7.**
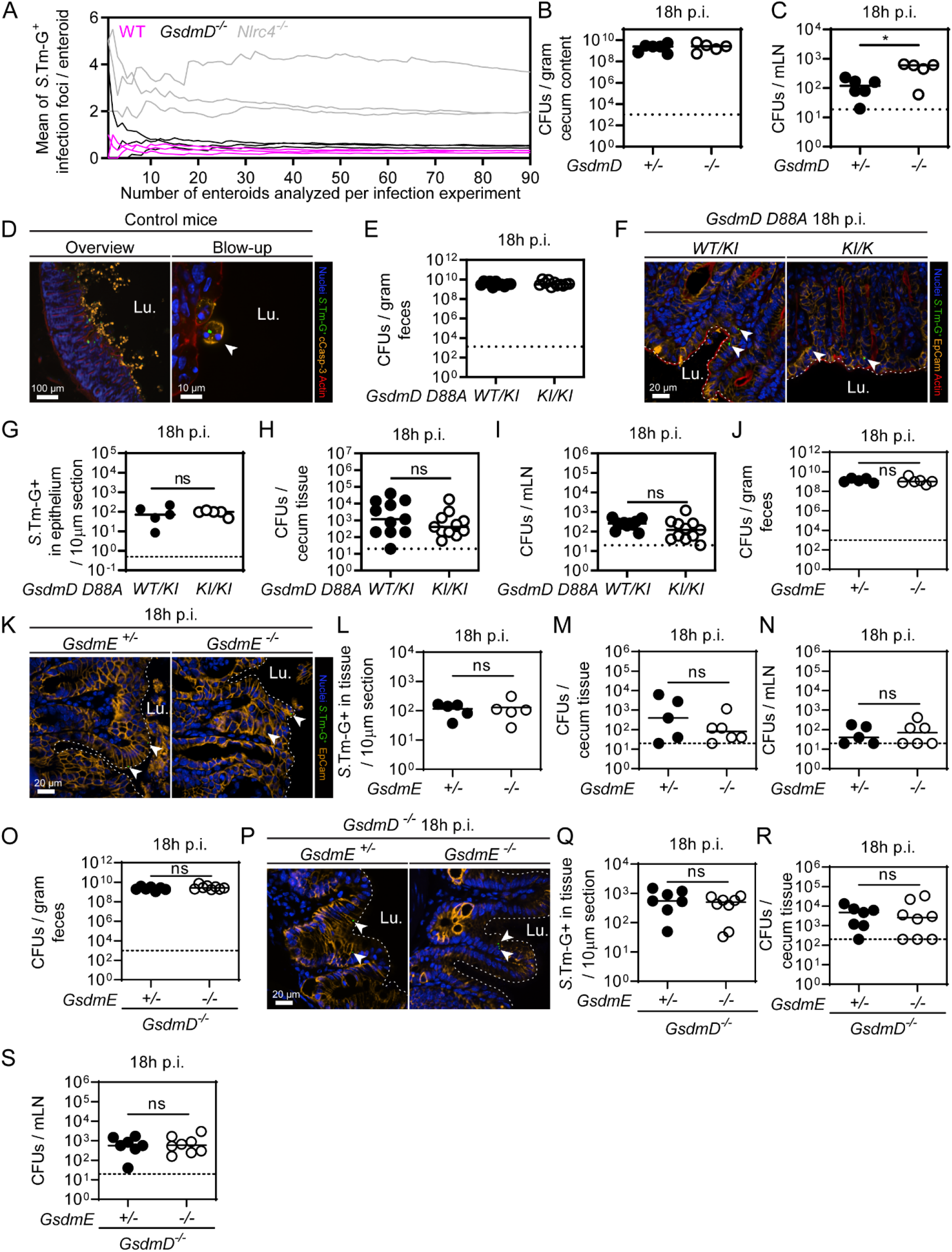
(Supplementary figure to Fig 4) (**A**) Determination of a sufficient sampling size for Fig 4A-B. Each line represents one replicate infection. 3 replicate infections per genotype. (**B-C**) 18h infection of *GsdmD*^*+/-*^ and *GsdmD*^*-/-*^ littermates with *S*.Tm harboring a *pssaG-GFP* reporter. *S*.Tm pathogen loads in (**B**) cecum content, and (**C**) mesenteric lymph nodes. (**D**) Representative micrographs of cecum tissue sections from a control mouse at 9h p.i. infected with *S*.Tm harboring a *pssaG-GFP* reporter and stained for cleaved Caspase-3. Arrowhead indicates an infected expelling enterocyte. Lu. - Lumen. (**E-I**) 18h infection of *GsdmD*^*WT/D88A*^ (referred as *WT/KI*) and *GsdmD*^*D88A/D88A*^ (referred as *KI/KI*) littermates with *S*.Tm harboring a *pssaG-GFP* reporter. (**E**) *S*.Tm pathogen loads in feces. (**F**) Representative micrographs of cecum tissue sections. Arrowheads indicate *S*.Tm-G^+^ in epithelium. Lu. - Lumen. (**G**) Microscopy-based quantification of *S*.Tm-G^+^ in epithelium. *S*.Tm pathogen loads in (**H**) cecum tissue, and (**I**) mesenteric lymph nodes. (**J-N**) 18h infection of *GsdmE*^*+/-*^ and *GsdmE*^*-/-*^ littermates infected with *S*.Tm harboring a *pssaG-GFP* reporter. (**J**) *S*.Tm pathogen loads in feces. (**K**) Representative micrographs of cecum tissue sections. Arrowheads indicate *S*.Tm-G^+^. Lu. - Lumen. (**L**) Microscopy-based quantification of *S*.Tm-G^+^ in cecum tissue. *S*.Tm pathogen loads in (**M**) cecum tissue, and (**N**) mesenteric lymph nodes. (**O-S**) 18h infection of *GsdmD*^*-/-*^*xGsdmE*^*+/-*^ and *GsdmD*^*-/-*^*xGsdmE*^*-/-*^ littermates infected with *S*.Tm harboring a *pssaG-GFP* reporter. (**O**) *S*.Tm pathogen loads in feces. (**P**) Representative micrographs of cecum tissue sections. Arrowheads indicate *S*.Tm-G^+^. Lu. - Lumen. (**Q**) Microscopy-based quantification of *S*.Tm-G^+^ in cecum tissue. *S*.Tm pathogen loads in (**R**) cecum tissue, and (**S**) mesenteric lymph nodes. In A, combined results from 2 independent experiments. In B, C, E, G-J, L-O, Q-S each data point represents one mouse. ≥5 mice per group from ≥2 independent experiments for each comparison. Line at median. Dotted line represents detection limit. Mann-Whitney U-test (ns - not significant, *p<0.05).

